# Are allocentric spatial reference frames compatible with theories of Enactivism?

**DOI:** 10.1101/170316

**Authors:** Sabine U. König, Caspar Goeke, Tobias Meilinger, Peter König

## Abstract

Theories of Enactivism propose an action-oriented approach to understand human cognition. So far, however, empirical evidence supporting these theories has been sparse. Here, we investigate whether spatial navigation based on allocentric reference frames that are independent of the observer’s physical body can be understood within an action-oriented approach. Therefore, we performed three experiments testing the knowledge of the absolute orientation of houses and streets towards north, the relative orientation of two houses and two streets, respectively, and the location of houses towards each other in a pointing task. Our results demonstrate that under time pressure, the relative orientation of two houses can be retrieved more accurately than the absolute orientation of single houses. With infinite time for cognitive reasoning, the performance of the task using house stimuli increased greatly for the absolute orientation and surpassed the slightly improved performance in the relative orientation task. In contrast, with streets as stimuli participants performed under time pressure better in the absolute orientation task. Overall, pointing from one house to another house yielded the best performance. This suggests, firstly, that orientation and location information about houses are primarily coded in house-to-house relations, whereas cardinal information is deduced via cognitive reasoning. Secondly, orientation information for streets is preferentially coded in absolute orientations. Thus, our results suggest that spatial information about house and street orientation is coded differently and that house orientation and location is primarily learned in an action-oriented way, which is in line with an enactive framework for human cognition.

## Introduction

Theories of embodied cognition understand cognition as being derived from the body’s interactions with the world (Varela et al. 1991; Wilson 2002). The enactive approach within embodied cognition theories emphasizes the importance of action, stating that even perception is an action (Noë 2004; O’Regan and Noe 2001). These theories assume that cognition is an embodied activity that includes the mind, the body, and the environment, and they stress the importance of action (Engel et al. 2013). Hence, embodied and enactive approaches to cognition provide modern frameworks to address current topics in cognitive science (Maye and Engel 2013).

Can we find empirical evidence for the enactive approach in the area of spatial cognition, a field of research that is naturally focused on interaction within the environment? A central aspect of spatial cognition is the differentiation of human navigation strategies based on two broad classes of reference frames: egocentric and allocentric reference frames (Burgess 2006; Gramann 2013; Klatzky 1998; Mou et al. 2004). Based on the work of Klatzky (1998), the egocentric reference frame is defined as relating the environment to the observer’s body, its physical position and orientation. Somatosensory information is gathered while moving in the environment. As the sensors are part of the body and move with the body, somatosensory information is initially coded within an egocentric reference frame. Spatial updating thus involves changes in egocentrically coded sensory information (Riecke et al. 2007) and is linked to self-motion (Simons and Wang 1998; Wang and Spelke 2000). In contrast, allocentric reference frames are independent of the observer’s physical body (Klatzky 1998). They are defined as being based on cardinal directions, geometric features, or environmental cues such as objects. Thus, spatial cognitive processes based on egocentric reference frames are naturally understood within the framework of enacted cognition, but further investigation is needed to determine whether navigation based on allocentric reference frames can also be understood within an action-oriented approach.

Many theories of spatial learning suggest that egocentric and allocentric reference frames develop together (Haith and Benson 1998; Nardini et al. 2006; Piaget and Inhelder 1967; Siegel and White 1975) and are used in parallel and interact (Burgess 2006; Gramann 2013; Ishikawa and Montello 2006; Waller and Hodgson 2006). Previous studies have shown, for example, that within an egocentric reference frame, viewpoint-dependent scene recognition (Diwadkar and McNamara 1997) and pointing accuracy (Shelton and McNamara 1997) improved when the tested viewpoint was aligned with the viewpoint from which the scene was learned. Other researchers have found that performance improves when imagined self-orientation was aligned to the observer’s own body orientation (e.g. Kelly et al. 2007). Within an allocentric reference frame, Mou and McNamara (2002) found an improved pointing accuracy from imagined viewpoints when the observer’s viewpoint was aligned with an intrinsic axis of an object array that represented aspects of the surrounding environment. They further investigated how locations of objects in a new environment are learned and stored in memory, and they introduced an intrinsic reference frame depicting object-to-object spatial relations combining aspects of an allocentric and egocentric reference system (Mou and McNamara 2002). Intrinsic and egocentric cues compete for the selection of reference frames in long-term memory, with a clear dominance of egocentric cues (Greenauer and Waller 2008; Street and Wang 2014). Remembered allocentric spatial relations guide egocentric action in space, which is updated while actively moving in the environment (McNamara 2002). Pointing accuracy was improved by aligning objects with the salient aspects of a large-scale environment (a salient landmark or the walls of a central building), which aspects were learned by actively walking in the tested environment (McNamara et al. 2003). Thus, information related to an allocentric reference frame is combined with egocentrically coded information to be translated into spatial action.

Spatial action (e.g. navigating from one location to another in a complex environment like one’s hometown) is crucial for everyday life. But how is this action achieved and the related knowledge stored in memory? It has been shown that while actively walking from one location to another, people develop “route knowledge” and sometimes also “survey knowledge” (Ishikawa and Montello 2006; Mallot and Basten 2009; Siegel and White 1975; Wiener et al. 2009). While the two locations are within a single vista space (Kelly and McNamara 2008; Waller and Hodgson 2006; Wang and Spelke 2000)—that is, a space that can be observed from one viewpoint—they are represented in one local reference frame, often following the orientation of a street in the neighborhood (Meilinger 2008a; Meilinger et al. 2014, 2016). When two locations are at a greater distance and can not be overlooked from a single viewpoint, called the environmental space (Montello 1993), multiple local reference frames that are learned by active navigation are combined. Thus, to move from one location to another in an environmental space, either, a network of pairwise combinations of local reference frames, graph knowledge, is formed (Meilinger 2008b) or multiple local reference frames are integrated into a global reference frame (McNamara et al. 2008; Poucet 1993). In navigating an environmental space, local and global reference frames may also be combined (Meilinger et al. 2014).

A global reference frame of a highly familiar space, such as one’s hometown, is supposed by some researchers to be stored in memory independent of orientation (e.g. Byrne et al. 2007; Sholl 1987). By contrast, others have suggested that familiar spaces are represented in one oriented global reference frame (e.g. McNamara et al. 2008; O’Keefe 1991; Trullier Wiener et al. 1997). Representing an environmental space in a global reference frame enables people to form “survey knowledge” and thus store distances and directions of locations, allowing them to take shortcuts and point from one location to another, invisible location. Alternatively, navigators rely on graphs consisting of local spaces and their metric interrelations to derive survey estimates (Chrastil and Warren 2014; Meilinger 2008a). Reference frames that are derived from subjective experience are supposed to have individual orientations. Instead, Frankenstein et al. (Frankenstein et al. 2012) found that spatial orientation in a familiar environmental space was best with the participants facing north. They argued that having city maps in Western countries that are oriented to the north facilitates performance in a pointing task when participants are aligned with this cardinal direction. Hence, reference frame orientation within a familiar environmental space can be based on individual experience or on the knowledge of cardinal directions.

But it is not only the navigating agent that determines how spatial representations are memorized and used. Gibson (1977) put forward the theory of affordances, proposing that objects and environments contain specific action possibilities, called affordances. These affordances require a relationship between the object or environment and the agent. On the basis of this relationship, object and agent then can work together. “When we perceive the object, we perceive the object’s affordances.”(Gibson 1979). Recently, it has been shown that action affordances, which indicate where one can go, are automatically extracted from a spatial scene (e.g. a room), and this results in the strong activation of a special visual area, the occipital place area (Bonner and Epstein 2017). In the present context, a town is typically composed of houses and streets, which offer different options for action. A house has a specific fixed location and orientation, and it invites people to meet there or to enter. In contrast, a street is used to walk or drive along, and it connects locations (e.g. houses) to each other. This difference in the affordances of houses and streets might also result in a difference of coding in terms of spatial aspects.

Within a single global reference frame that is centered, for example, on cardinal directions, the location and orientation of each object (e.g. a house) could be coded separately in memory. An advantage of this strategy is efficiency, as the amount of information to be stored scales linearly with the number of objects. By contrast, gaining information for spatial behaviors (e.g. navigating from one house to another) requires access to the information related to the starting and ending locations and to the cognitive processes involved in combining this information. Thus, storage efficacy of this strategy inflicts a time penalty when accessing action-related information. Alternatively, in a memory structure derived from experience, the locations and orientations of objects could be primarily learned and stored in relation to each other. As a disadvantage, the amount of information to be stored scales supra-linearly (e.g. quadratically, when assuming all-to-all relations between objects) with the number of objects. However, retrieving information to enact a spatial behavior (e.g. navigating from one house to another, or pointing from one house to another) only requires direct access to the stored relation of starting and ending locations; no further cognitive considerations are necessary. Thus, this approach yields an advantage in speed at the expense of storage needs. Therefore, these two types of coding differ qualitatively in terms of storage needs and speed of access to action relevant information.

Following this reasoning, we investigated whether spatial knowledge based on allocentric reference frames is nevertheless coded and utilized in an action-oriented way. We selected stimuli in the form of houses and streets of a single familiar city and investigated how they relate to different types of actions and how they might thus be coded in different styles. In order to investigate the two types of coding spatial information and avoid naively equating the specific experiments with forms of spatial reference frames, for the purpose of the present study we call the two types of coding *unitary code* and *binary code*. A *unitary code* relates to a single object in terms of its orientation with respect to cardinal directions, whereas a *binary code* relates to a combination of objects, thus representing the orientations of objects in terms of object-to-object relations. In view of the advantages and disadvantages of each, it is conceivable that humans may use either a *unitary* or *binary coding* strategy, or both.

In a variety of spatial navigation tasks, men showed faster reactions and better performance than women (Masters and Sanders 1993; Moffat et al. 1998; Newhouse et al. 2007; Woolley et al. 2010). These results point to systematic differences between the genders and, therefore, we analyzed gender differences in our tasks. However, these gender differences do not obviously relate to the specific question of *unitary* versus *binary coding*. Thus, we did not expect systematic differences in relation to our research questions (i.e. no interactions between gender and tasks), but rather a confirmation of men’s better performance (i.e. solely a main effect of gender).

We performed three main experiments to address the coding of spatial information for single objects and for object-to-object relations. The first experiment tested knowledge of absolute orientation (relative to north), thus investigating *unitary coding*. We realized this with two different stimuli types: frontal views of houses as localized stimuli and views along streets as the means to navigate from one location to another. The second experiment tested knowledge of relative orientation, thus investigating *binary coding*. We achieved this by using the same stimuli that were used in the first experiment. That is, we queried the relative orientation of frontal views of two houses as well as the relative orientation of two views along streets. The third experiment tested navigation knowledge in the form of a more classical pointing task. This task used houses as the sole stimulus, and asked participants to point from one house to another. Please note that here our primary focus was on allocentric reference frames and not on the influence of bodily alignment relative to the stimuli shown. We analyzed the influence of bodily position during the experiment only as a secondary aspect. To determine if participants could directly access the required spatial information or whether further cognitive processing steps were needed, participants either performed under time pressure or had unlimited time to respond (for further details, see the Methods section). Specifically, we tested the following hypotheses:

- H_0_: Spatial information about houses is coded separately for each house relative to a common reference frame—that is, the orientation of a house with respect to cardinal directions (*unitary code*). Accessing action-relevant relations between houses out of the *unitary* information requires time-consuming cognitive reasoning.
- H_1_: Spatial information about houses is coded in direct relation to other houses (*binary code*). This direct coding of house-to-house relations allows for the relations between houses to be accessed quickly and intuitively, making it suitable for the generation of behavior.
- H_2_: Spatial information related to stimuli such as streets is primarily coded in a *unitary code*, which is different from how houses are coded. Thus, the retrieval of directional information that is directly relevant to an action does not require time-consuming cognitive processing.

In summary, the present study tests the availability of *unitary* and *binary coding* of spatial knowledge for human behavior.

## Methods

### Participants

Overall, we recruited 69 participants who, previous to participating in our study, had lived at least one year in Osnabrück. Thirty-nine participants (24 females, mean age of 24.2 years, SD = 2.47 years) took part in the first and second experiments, which involved houses as stimuli and enforced a restricted response time. The remaining 30 participants (19 females, mean age of 23.3 years, SD = 2.34 years) performed the first and second experiment, which involved houses as stimuli and infinite response time as well as with street stimuli with restricted and infinite response time. This group additionally performed a third, pointing experiment, which involved houses as stimuli and enforced a restricted response time. A summary of the experiments can be found in Table 1, and the results are illustrated in Fig 1.

**Table 1:**
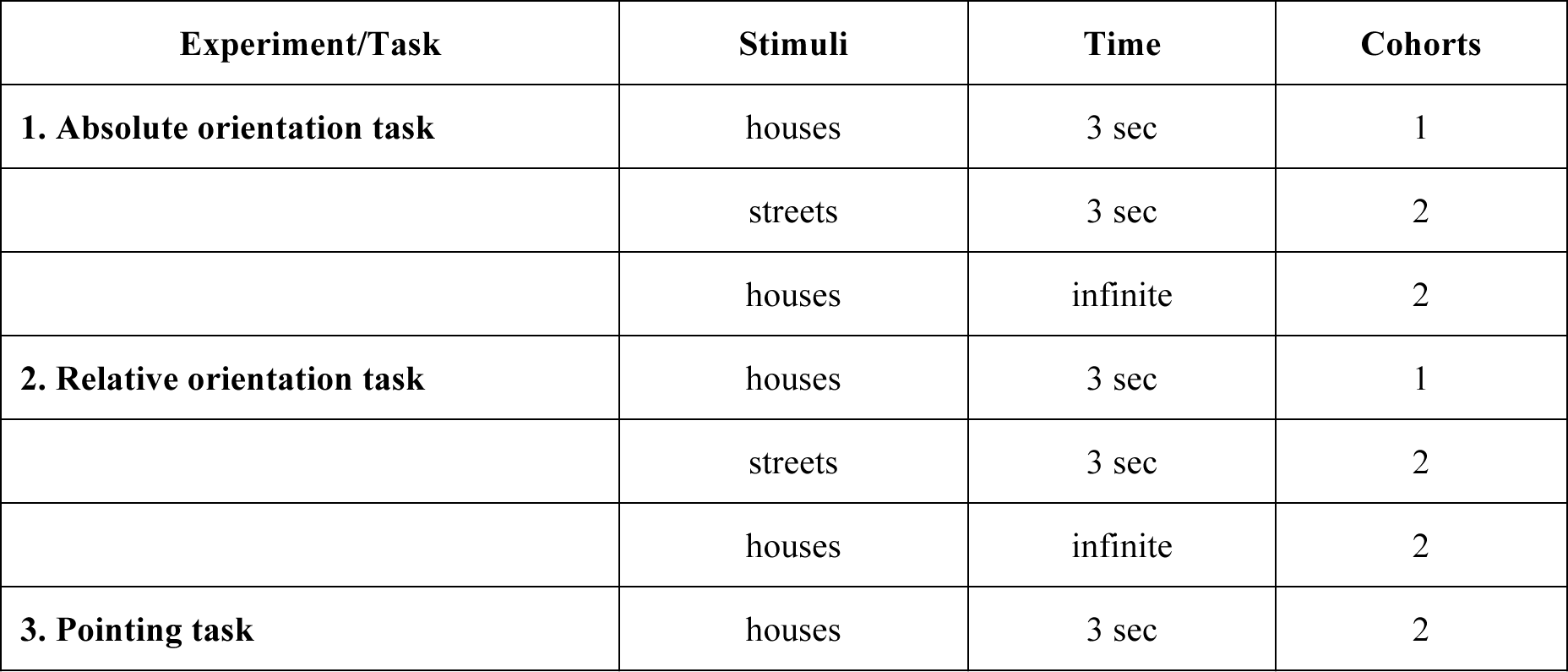
Overview of the complete experimental design.

| Experiment/Task | Stimuli | Time | Cohorts |
| --- | --- | --- | --- |
| **1. Absolute orientation task** | houses | 3 sec | 1 |
|  | streets | 3 sec | 2 |
|  | houses | infinite | 2 |
| **2. Relative orientation task** | houses | 3 sec | 1 |
|  | streets | 3 sec | 2 |
|  | houses | infinite | 2 |
| **3. Pointing task** | houses | 3 sec | 2 |

**Fig 1.**
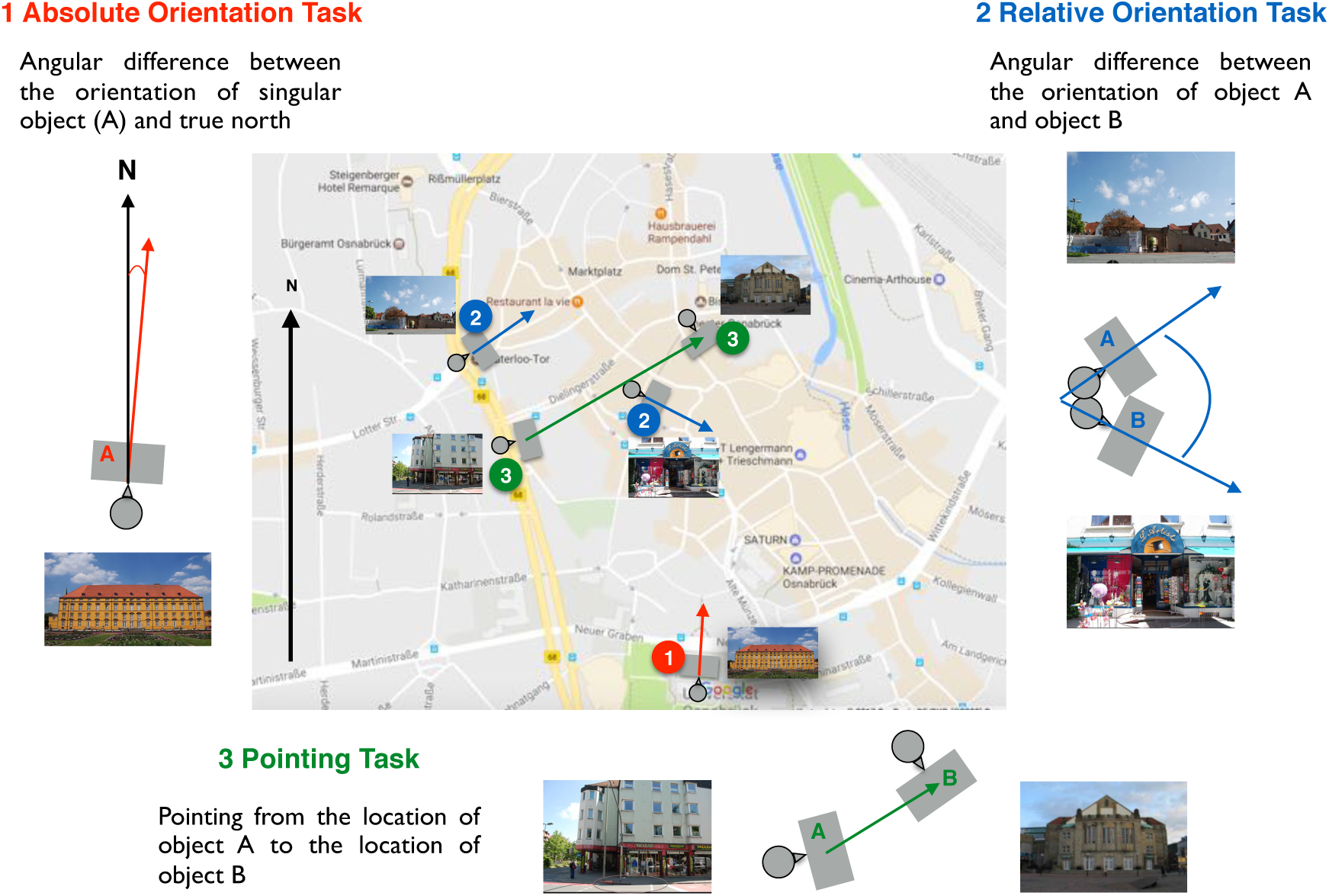
Overview of the experimental tasks. illustrated on a central part of the Osnabrück city map (OpenStreetMap) and in a schema; (1 = Absolute Orientation Task, 2 = Relative Orientation Task, 3 = Pointing Task). For further explanation, see main text.

### Training

Each participant performed a response training, in which they trained first to respond within 3 seconds and then learned how to read the directional arrow on the screen and what the required behavioral response was. In this training, one arrow surrounded by an ellipsoid appeared on the upper screen, and another appeared on the lower screen, each pointing in different directions. The participants had to compare the two arrows and then select the arrow that pointed more straight upward by pressing the “up” button to select the arrow on the upper screen or pressing the “down” button to select the arrow on the lower screen. On each trial, they got feedback as to whether they decided correctly (green frame), incorrectly (red frame), or failed to respond in time (blue frame). In order to finish the training, the participants had to respond correctly without misses in 48 of 50 trials (95%). This procedure ensured that the mechanisms of the experiment were well understood and did not present any difficulties, even under time pressure.

## Experimental Design

### 1. Absolute orientation task

In the absolute orientation task, we assessed the participants’ ability to estimate the orientation of single houses in relation to the north cardinal direction in a two-alternative forced-choice task (see Fig 1, the red marker “1” and the red arrows, and Fig 2). As stimuli, we showed real-world front-view photographs of houses in the city of Osnabrück (Fig 2a). For each of the 156 trials, we presented participants with two separate screens, one above the other, both showing the same image. On both images, an arrow within an ellipsoid depicting a compass was shown. One of the arrows pointed to the north cardinal direction, whereas the other arrow pointed in a direction that diverged from north by some amount between 0 and 330° (in steps of 30°). The orientation of the arrow in the second stimulus was random. The resulting small fraction of ambiguous trials was not considered in the data analysis. The participants had to choose in which of the images the arrow pointed correctly towards north by pressing either the “up” (upper screen) or “down” (lower screen) button on a response box. To test the influence of response time, we designed the absolute orientation task using photographs of houses as stimuli and either to enforce a restricted response time (3 s) or allow infinite time for response (Table 1). To test the influence of different kinds of stimuli, participants also performed the absolute orientation task with photographs of streets serving as stimuli (Fig 2b) in the 3-second response mode.

**Fig 2.**
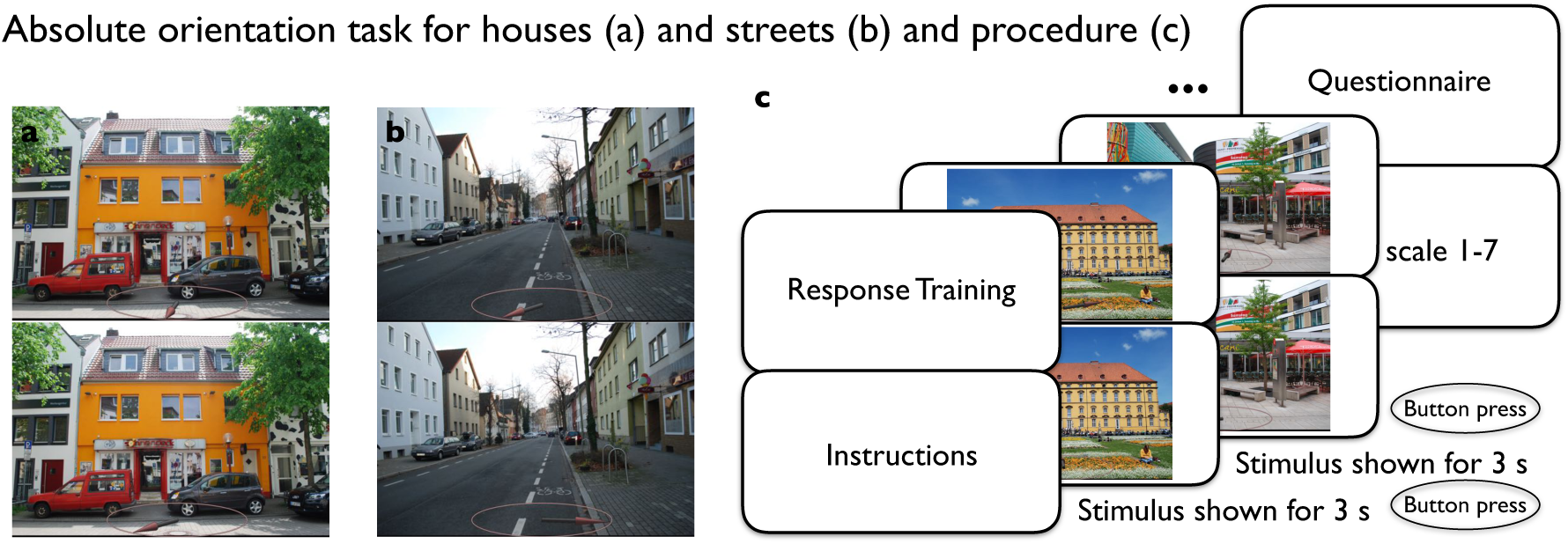
Absolute orientation task. The illustration shows one sample trial of the absolute orientation task using house stimuli (a) and street stimuli (b), respectively. An example of the procedure of the absolute orientation task can be seen in (c). An image was shown on two monitors, one above the other, for 3 seconds. The images were overlaid with arrows within an ellipsoid, depicting a compass. In each example, one arrow points correctly to north and the other to a different direction (diverging from north by some amount between 0 and 330° in steps of 30°). Participants had to choose the arrow that correctly pointed north.

### 2. Relative orientation task

In the relative orientation task, we assessed the ability to estimate the orientation of houses relative to each other (see Fig 1, blue marker “2” and the blue arrows, and Fig 3a). We designed a two-alternative forced-choice task using the same frontal-view photographs from the Osnabrück area as those used in experiment 1. Each of the 72 trials in the relative orientation task consisted of a fixed triplet of images: one priming image and two target images (see stimulus preparation). In each trial, first the priming picture was shown on both screens for 5 seconds. After the priming stimulus was turned off, the two target stimuli appeared, one on the upper screen and the other on the lower screen. The task of the participants was to select the upper or lower target image as being more closely aligned with the orientation of the priming image by pressing either the “up” or “down” button on a response box, respectively. Again, house stimuli were tested with both a restricted response time (3 s) and infinite response time (Table 1). The relative orientation task was also performed with photographs of streets as stimuli (Fig 3b) with a 3-second response time.

**Fig 3.**
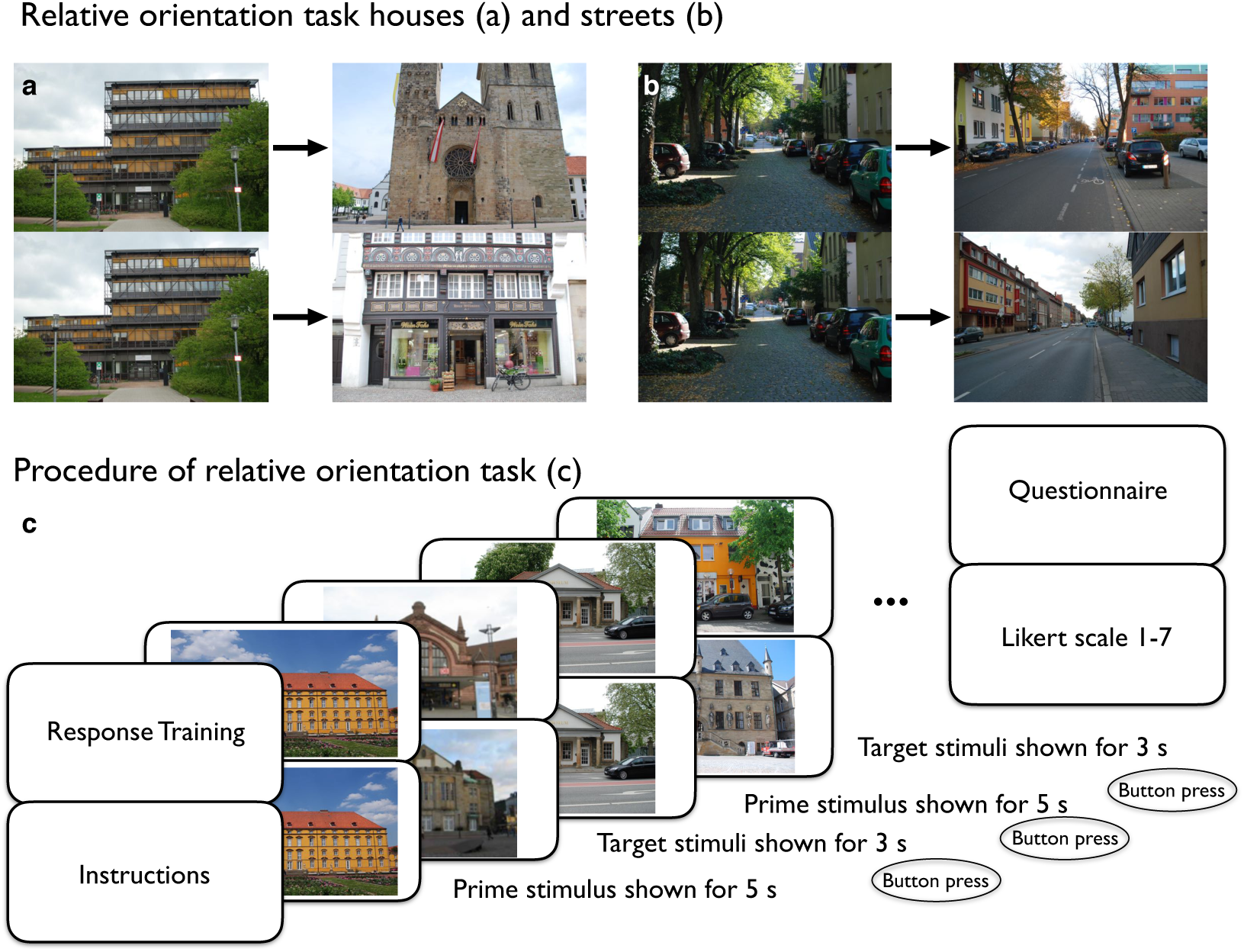
Relative orientation task. The illustration shows one sample trial of the relative orientation task using house stimuli (a) and street stimuli (b). An example of the procedure of the relative orientation task can be seen in (c). First, a priming image (left) was shown for five seconds. Then, two different target images appeared on the two screens, one above the other (right). One of the target images has the same orientation as the priming image, while the other differs by some amount between 0 and 330° in steps of 30°. The task of the participants was to select the image that was oriented in the same direction as the priming image.

### 3. Pointing Task

Participants performed a pointing task, which is an established paradigm in spatial navigation studies, to assess the knowledge of spatial relations between the locations of tested objects. We used the same stimulus material of houses (only) and selected pairs of images (priming and target images; see Fig 1, green marker “3” and the green arrows, and Fig 4). For each of the 144 trials, the priming stimulus appeared (on both screens) for 5 seconds, and afterward the target stimulus appeared (again, on both screens). Similar to the absolute orientation task, we overlaid each target image with an arrow within an ellipsoid, this time depicting the pointing direction. One of the arrows was correctly pointing in the direction of the priming image location, and the other arrow was pointing in a random direction with varying degrees (0–330° in 30° steps). In a two-alternative forced-choice task, the participants had to choose the image on which the arrow pointed correctly towards the priming stimulus by pressing either the “up” or “down” key.

**Fig 4.**
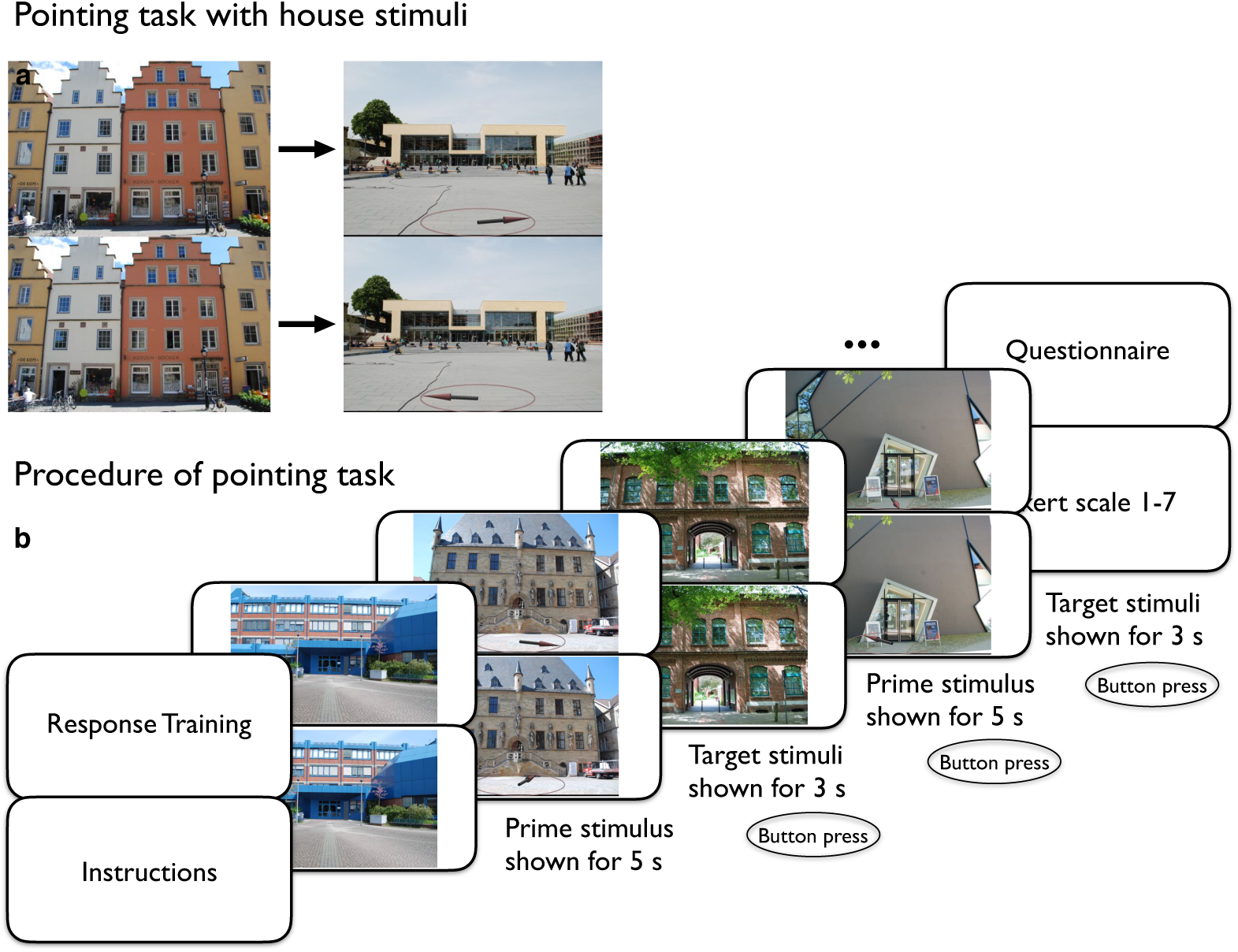
Pointing task with house stimuli. The illustration shows one sample trial of the pointing task using house stimuli (a) and the procedure of the pointing task (b). First, a priming image was shown for 5 seconds (left) on both monitors. Then, a target image was shown for 3 seconds (right), the same on both monitors. One of the target images was overlaid with an arrow pointing correctly towards the priming image, whereas the other differed by some amount between 0 and 330° in steps of 30°. The participant had to choose the arrow that pointed correctly towards the priming image

#### Stimulus Recordings and Preparation

Overall, we used two different stimulus sets: real-world photographs of houses and of streets taken in the Osnabrück city area. All photographs were taken with the same type of camera and used the same camera settings. The distance to the objects was set between 5 and 15 meters, depending on the viewpoint that resulted in the most intuitive perspective. That is, that the whole house or street was visible, with little else in the image that could serve as a distractions. While taking the photos, we ensured good lighting conditions and avoided occlusions of the main motif. All photos were taken frontally, such that for houses, the viewing direction of the photo was aligned with the main facade of the house (see Figs 1–4). Regarding streets, we only picked straight segments so that the viewing perspective followed the course of the street (see Figs 2 and 3). The recording of the cardinal direction was conducted using a mobile phone at the same time the pictures were taken. To improve reliability, we repeated these measurements five to seven times and calculated the average for each photograph. For the relative orientation task and the pointing task, we selected fixed sets (pairs and triplets) of images that were presented together within one trial. The selection of the priming images for both tasks was based on a pilot study (8 participants) in which all participants rated the familiarity of all stimuli. The most familiar images were chosen as priming images. Additionally, for the relative orientation task, we applied an optimization algorithm that enabled us to select a target image whose cardinal direction was very closely aligned to the orientation of the priming image (<10°) as the correct target image. By contrast, the orientation of the incorrect target image randomly varied compared to the correct target image in roughly equal steps of 30°.

#### Procedure

During the experiments, all participants sat approximately 60 cm from a six-screen monitor setup in a 2x3 arrangement. The images were only presented on the two central screens (one above the other), while the other four monitors on either side were switched off. The resolution of the original images was 3872x2592 pixels, but all photographs were rescaled so that each image was presented in full screen (2160x1920 pixels) on one of the monitors. Each participant started with the response-training task. Written instructions were given on the screen. Participants continued by pressing a button on the control box. The order of the experiments was balanced across participants. The order of stimulus presentation in each experiment was also randomized so that each participant was presented with a different sequence of stimuli. In all tasks, written instructions were followed by a stimulus set on the two central screens. In the absolute orientation, the stimulus set consisted of the same stimulus on the upper and lower screen, both of which were overlaid with arrows in an ellipsoid (see above). After the press of a button, another stimulus set appeared on the screens. All in all, the procedure was repeated with 156 stimuli. In the relative orientation and pointing task, first a priming stimulus appeared on both screens and was followed by different target stimuli on the two screens (see above). Participants selected the correct stimulus by pressing the “up” or “down” control buttons. This sequence was repeated 72 times. In the pointing task, the priming stimulus was followed by a target stimulus that was depicted on both screens overlaid with arrows in an ellipsoid. This task included 144 trials. At the end of the experiment, all participants rated their familiarity with the seen stimuli. The latter data are not presented within this study. All participants were informed about the experiment and filled out the required consent form. Ethical IRB approval was obtained prior to the experiment from the ethics committee of the University of Osnabrück. The participants were either reimbursed with 7 Euros per hour or earned an equal amount of “participant hours,” which are a requirement in most students’ study programs. Overall, the experimental session in cohort 1 took about 1 hour, and the participants in cohort 2 took about two and a half hours.

## Results

### Performance in the absolute and relative orientation tasks with house stimuli and restricted as well as infinite response times

First, we compared the performance in the absolute orientation task, measuring knowledge of cardinal orientation and the relative orientation task, evaluating houses as stimuli. Hence, we calculated the fraction of correct responses separately for each participant and task. Then, we performed a three-way mixed-measure ANOVA with performance as the dependent variable, task as a repeated factor (absolute vs. relative), and response mode (3 s vs. infinite) and gender as between-participant factors. As described above, we decided to include gender in order to account for previously reported gender differences in the domain of spatial cognition. The test revealed no main effect of task (F(1,65) = 2.648, p = .109, partial η^2^ = .039). However, we observed a significant main effect for the response mode (F(1,65) = 56.479, p < .0001, partial η^2^ = .465), as well as a main effect of gender (F(1,65) = 20.968, p < .0001, partial η^2^ = .244). Post hoc comparisons with Bonferroni corrections confirmed that infinite response time improved performance significantly compared to the 3-second response time (p < .001). Additionally, in this test, males outperformed females (p < .001). There were no interactions of gender with response time (F(1,65) = .083, p = .774, partial η^2^ = .001) or task (F(1,65) = .585, p < .447, partial η^2^ = .009). The three-way interaction of all factors was also not significant (F(1,65) = .320 p =.859, partial η^2^ < .001). However, we found an additional significant interaction between task and response mode (F(1,65) = 17.630, p < .0001, partial η^2^ = .213). As a follow-up, we conducted two separate paired t-tests (one for each participant group) to investigate potential differences between the absolute and the relative orientation tasks, one for each of the two reponse modes. The Bonferroni corrected test statistics revealed opposing effects: With a 3-second response time, participants performed significantly better in the relative orientation task (M = 56.04, SD = 6.38) than in the absolute orientation task (M = 52.84, SD = 6.39, *t* (38) = −2,418, *p* = .021). With infinite response time, participants showed a significantly better performance in the absolute orientation task (M = 66.85, SD = 8.88) compared to the relative task (M = 60.55, SD = 7.48), *t* (29) = 3,610, *p* < .001; see Fig. 5). Thus, men outperformed women in both tasks, though there was no interaction between gender and task. Further, we conclude that, in agreement with Hypothesis 1, our participants had better access to information about house-to-house orientation than about the cardinal orientation of houses. However, given enough time for cognitive reasoning, performance improved strongly for the absolute orientation task, whereas performance in judging house-to-house orientation only improved moderately.

**Fig 5.**
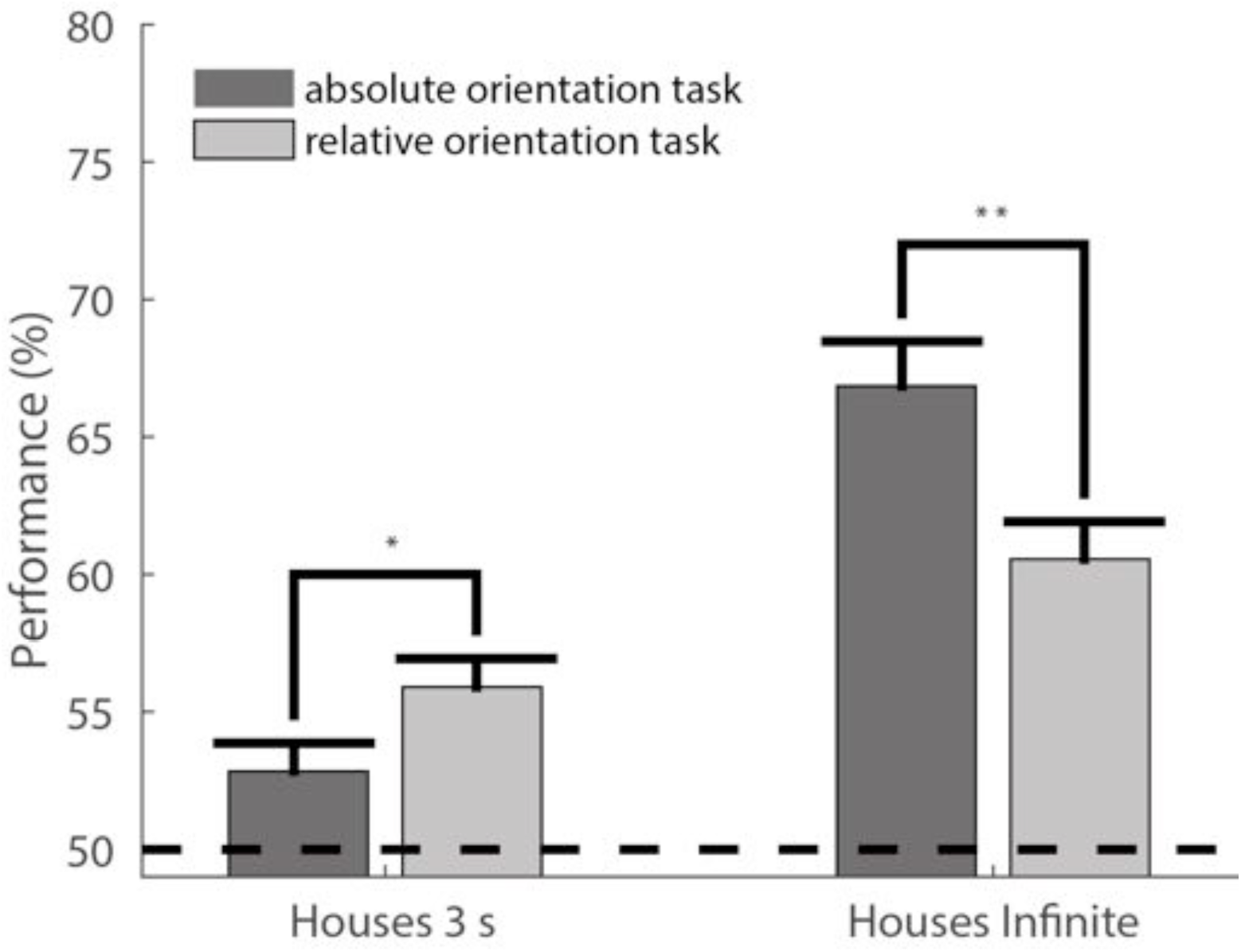
Performance of absolute and relative orientation task with restricted and infinite response modes. The two bars on the left show the performances in the 3-second response mode with house stimuli, whereas the two bars on the right show the performance levels for the infinite response mode with house stimuli. The black dashed line indicates the chance level (50%). The error bars indicate the standard errors of the mean (SEM), and the asterisks indicate significance at the respective thresholds of ^∗^p < 0.05 and ^∗∗^p < 0.01. The y-axis displays the mean performance levels for the different experiments. The dark grey bars refer to the absolute orientation task, whereas the light grey bars refer to the relative orientation task.

### Performance in the absolute and relative orientation tasks with house and street stimuli

In order to compare performance between both stimuli sets, we performed a mixed-measure ANOVA with test performance as the dependent variable, task (relative vs. absolute) as a repeated factor, and stimulus set (houses vs. streets) and gender as between-participant factors. Neither the task (F(1,65) = .153, p = .697, partial η^2^ = .002) nor the stimulus set (F(1,65) < .001, p = .991, partial η^2^ < .001) revealed a significant main effect. Thus, the absolute orientation task and the relative orientation task were of comparable difficulty, and the demands of estimating the orientation of houses and streets were comparable as well. The factor of gender revealed a significant main effect (F(1,65) = 18.454, p < .0001, partial η^2^ = .221), but none of its interactions were significant. The three-way interaction was not significant (F(1,65) = .549, p = .461, partial η^2^ = .008). Importantly, however, the interaction of task and stimulus set was significant (F(1,65) = 9.105, p = .004, partial η^2^ = .123). To investigate this further, we contrasted the absoute and the relative orientation tasks seperately for both stimuli sets (Fig 6). As shown before, in the restricted time condition, participants jugding the house stimuli performed significantly better in the relative orientation task (M = 56.04, SD = 6.38) than in the absolute orientation task (M = 52.84, SD = 6.39, *t* (38) = −2,418, *p* = .021). By contrast, participants jugding street stimuli performed significantly better under the restricted time condition in the absolute orientation task (M = 56.51, SD = 6.10) compared to the relative orientation task (M = 52.48, SD = 7.01, *t*(29) = 2,414, *p* = .022). These results suggest that people access the orientation of houses and streets differently. Although the house-to-house orientation is more easily accessible than the absolute orientation of houses, the absolute orientation of streets is easier to judge than two streets’ relative orientation to each other.

**Fig 6.**
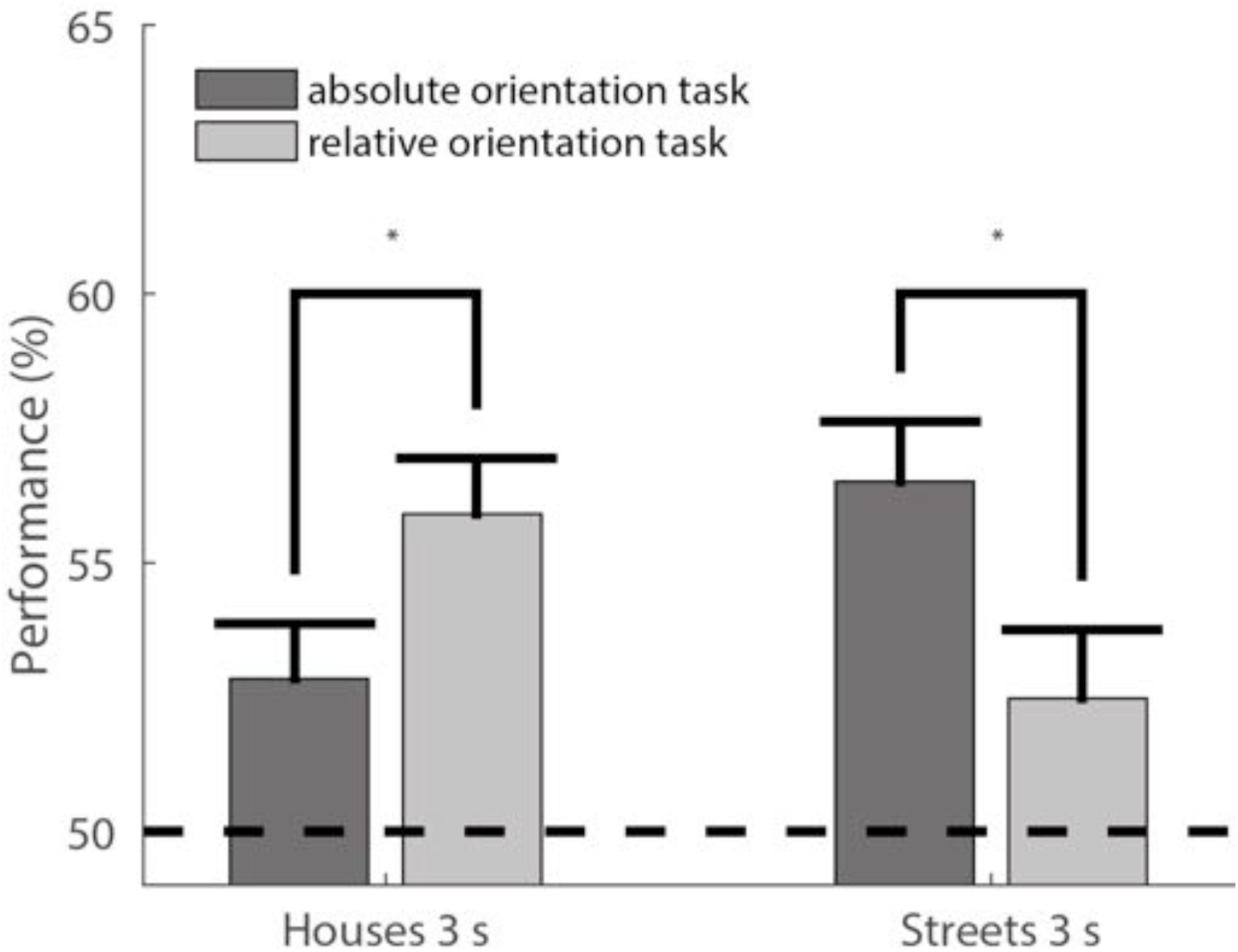
Performance in absolute and relative orientation tasks with house and street stimuli. The two bars on the left show the results of using house stimuli and the 3-second response mode, whereas the two bars on the right show the results of using the street stimuli and the 3-second response mode. The black dashed line indicates the chance level (50%). The error bars indicate SEM, and the asterisks indicate significance at the threshold of ^∗^p <0.05. The y-axis displays the mean performance levels for the different subtasks. The dark grey bars refer to the absolute orientation tasks, while the light grey bars refer to the relative orientation tasks

### Performance in the absolute orientation, relative orientation, and pointing tasks with house stimuli and restricted response time

We then compared performance in the pointing task to performance in the absolute and relative orientation tasks with house stimuli and 3-second response time. Please note that this comparison involves two different cohorts and thus includes a between-subject design. For statistical testing, we conducted two separate one-way ANOVAs, comparing the performance in the pointing task to the performance in the two orientation tasks. As shown in Fig 7, performance in the pointing task was significantly better than in the absolute orientation task (F(1,68) = 14.256, p < .0001, partial η^2^ = .175). However, the difference between the performances in the pointing and relative orientation tasks did not reach significance (F(1,68) = 3.621, p = .061, partial η^2^ = .051). Our results indicate that knowledge about relative object orientation and location are significantly better than the estimation of absolute object orientation to cardinal directions.

**Fig 7.**
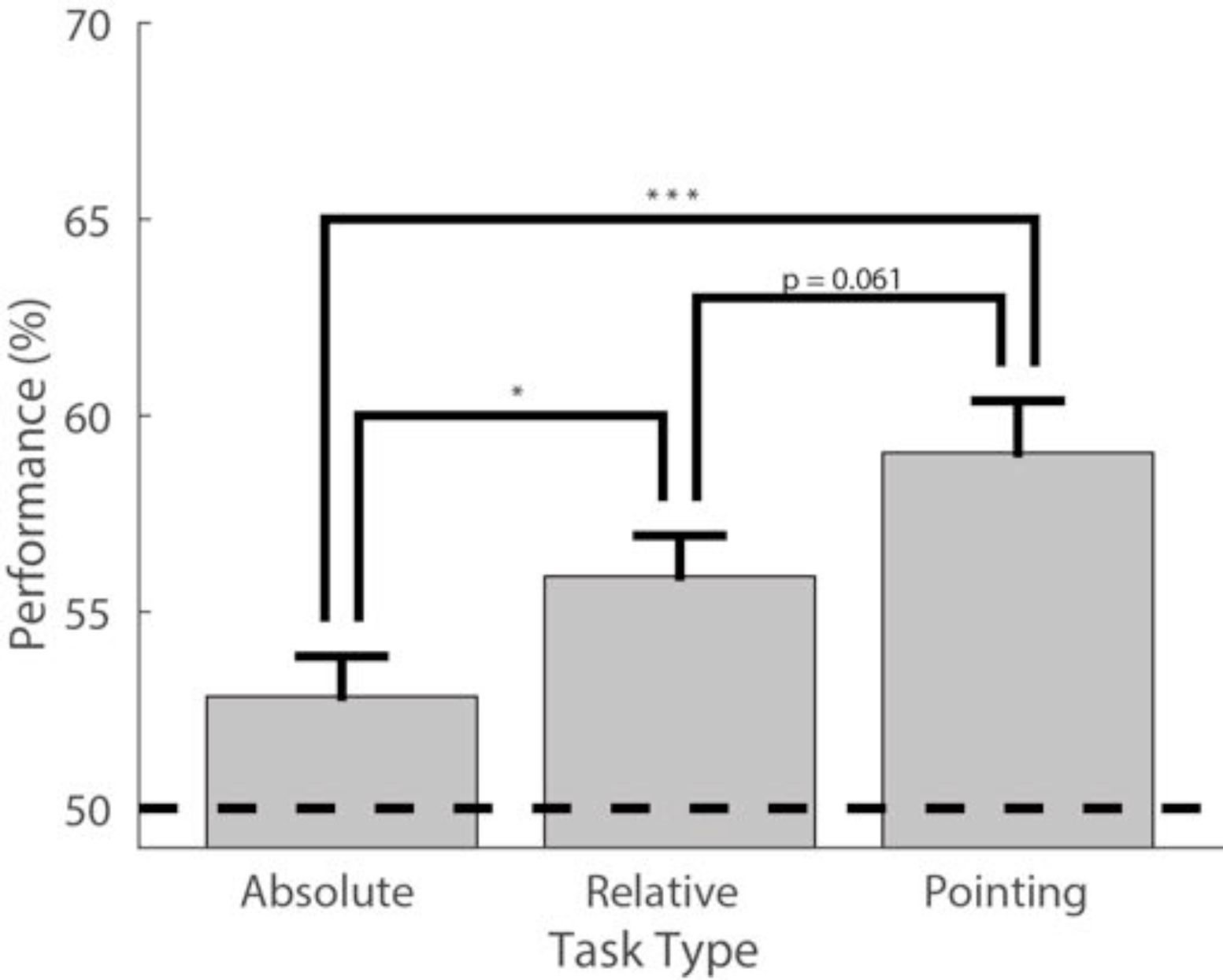
Performance in the absolute orientation, relative orientation, and pointing tasks. The y-axis displays the mean performance levels for the different experiments, using house stimuli and the 3-second response time. On the x-axis, the leftmost bar shows the performance of the absolute orientation task, the central bar the performance for the relative orientation task, and the rightmost bar the performance of the pointing task. The black dashed line indicates the chance level (50%). The error bars indicate SEM, and the asterisks indicate significance at the respective thresholds of ^∗^p < 0.05, ^∗∗^p < 0.01, and ^∗∗∗^p < 0.001.

## Distance

To estimate the influence of distance between the priming and target stimuli used in the relative orientation task, we first calculated the average distance of the two targets to the respective priming stimulus. With these data, we performed a linear regression of distance and performance (Fig 8). In the investigated range, up to 2.5 km, we did not find a significant relation between the average distance and the performance (R^2^ = 0.00658, p = 0.380). A more detailed analysis (e.g. taking the individual distances and the angle between the two prime target vectors into account) increases considerably the number of degrees of freedom and would require much more data. Thus, presently we refrain from making a conclusive statement regarding the dependence of performance on the geometry of primes and targets.

**Fig 8.**
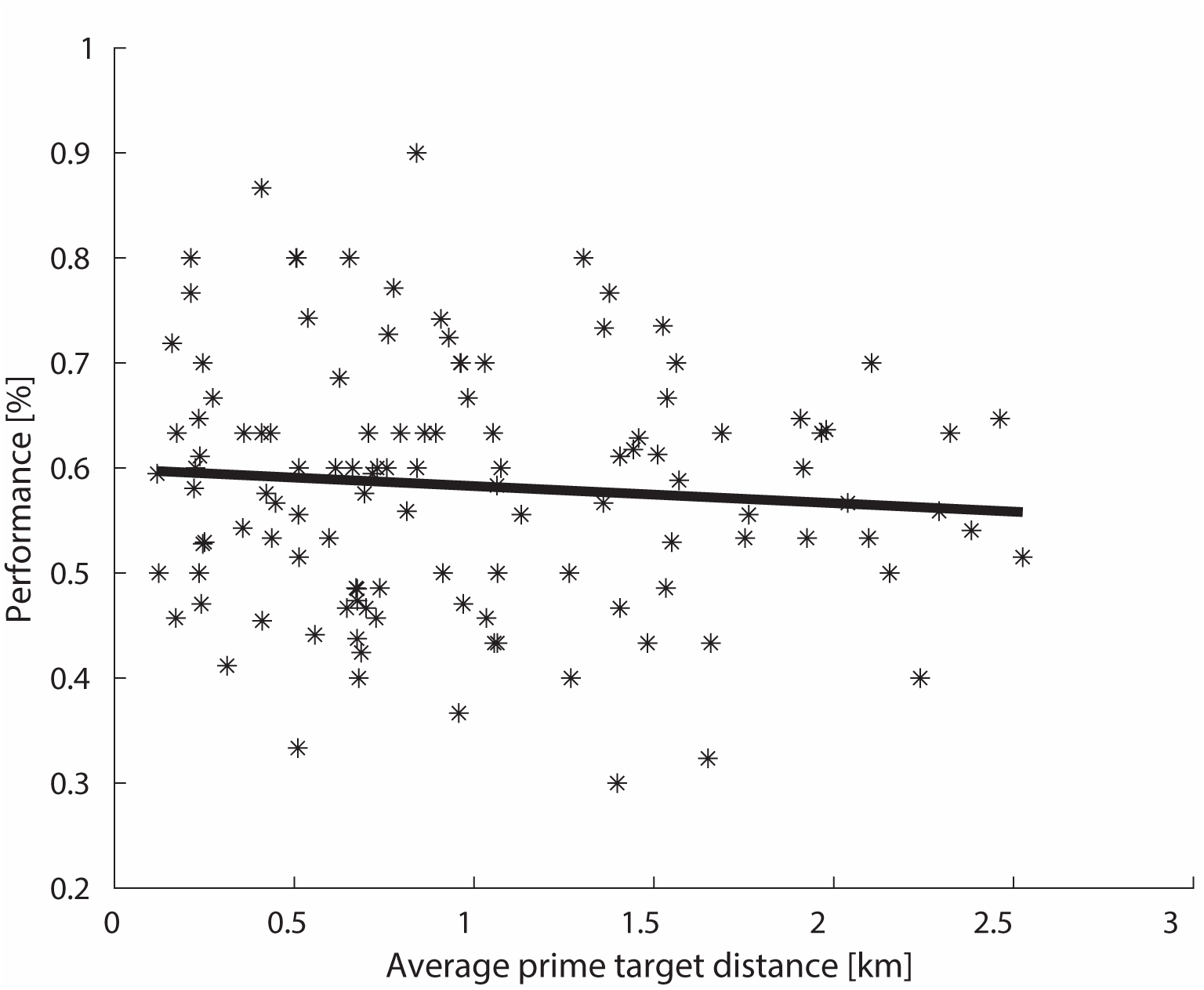
Influence of distance between priming and target stimuli on the overall performance. The asterisks show single data points, and the black line shows the linear regression for the distance between the performance and the average prime target distance.

### Angular Difference

The key question of our study asks how people learn and represent absolute and relative differences of orientation and location with respect to real-world houses and streets. In order to evaluate whether our introduced experimental manipulation, namely the quantitative difference in orientation between two real-world objects, had an influence on the participants’ performances, we grouped the stimuli accordingly in bins of 30°. We then applied a generalized linear model to all the data, separately for each of the different experiments (see Table 1). The participants’ binominal responses (i.e. correct, wrong) were fitted against the different levels of angular difference using a logit link function. Overall, the effect was rather weak, meaning that larger angular differences between two stimuli did not always (i.e. for each participant and condition) result in better performance. In particular, the performance in the two experimental conditions that had an overall near-chance performance (i.e. absolute orientation task with house stimuli with 3-second response time as well as relative orientation task with street stimuli with 3-second response time) had no demonstrable dependence on angular differences. As a result, individual psychometric evaluations could not be performed. However, in accordance with the previous findings, all other conditions that showed an overall reasonable performance (>55%) demonstrated a significant (p < 0.05) change in slope. Hence, we can conclude that on a broad-enough scale, larger angular differences between two choices led to better performance.

### Influence of Viewing Direction

Burte and colleagues suggested that the alignment of viewing direction and true cardinal direction can have a strong impact on spatial judgments (Burte and Hegarty 2012, 2014). Hence, we aimed to determine whether the deviation of the viewing direction from true north, as indicated by arrow direction, also influenced our participants’ decisions. Therefore, we first calculated the average performance for each stimulus, separately for the different experiments. Then, we analyzed the performance as a function of the difference between the viewing direction and true north. Specifically, we determined the direction that resulted in the best performance. To account for the circular nature of cardinal direction information, we then fitted a cosine to the data and as a result got the phase φ, which yielded the best fit and explained the variance of fit (R^2^). We performed this analysis for both the relative and absolute orientation tasks. We found no effects for any of the relative task conditions. Whereas, in the absolute task with restricted response time, the performance was clearly modulated by an alignment of the viewing direction and true north (Fig 8). The best fit for the absolute orientation task with house stimuli and restricted response time was achieved when they were oriented very close to the full circle with φ = 359.28°, reaching an R^2^ of 0.287. The best fit for the absolute orientation task with street stimuli and restricted response time had a φ = 9.52°, reaching an R^2^ of 0.301. In a second step, we tested for statistical significance, and consequently divided the data into two bins: In the first bin, we put all stimuli in which the correct arrow (north) pointed in the forward direction (0°+/-90°). In the second bin, we put the remaining stimuli with the correct arrow pointing backwards. Consequently, for the absolute orientation task one-way ANOVAs showed that with restricted response time performance was significantly better when the correct arrow pointed in the forward direction for house stimuli (F(1,139) = 36.930 p < .0001, partial η^2^ = .211) and street stimuli (F(1,142) = 57.337, p < .0001, partial η^2^ = .289). With an infinite amount of response time, the size of the effect decreased dramatically, yielding a less circular (R^2^ = 0.014) but still significant fit (F(1,140) = 4.065, p = .046, partial η^2^ = .028). In summary, in the absolute orientation task, participants performed best when the correct arrow (true north) was pointing right in front of them and was thus aligned with their own viewing orientation. Interestingly, this effect was most visible when decisions under time pressure (3-second response mode) were required. We found no effect of the viewing direction on performance in the relative orientation task.

**Fig 9.**
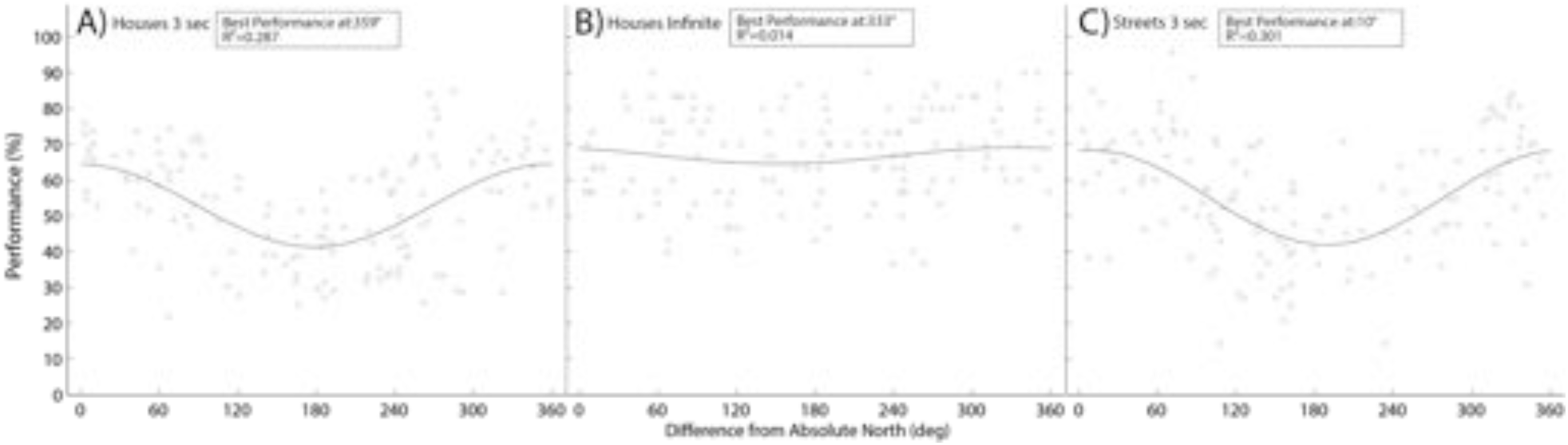
Influence of viewing direction in the absolute orientation tasks. The three panels show the influence of viewing direction in the absolute orientation tasks with different stimuli and response times. The left panel shows the data for the absolute orientation with houses and 3 second response time, the center panel shows the data for the the absolute orientation with houses and infinite response time, and the right panel shows the data for the absolute orientation with streets and 3 second response time. The y-axis displays the average performance for each image as a mean across participants. The x-axis indicates the difference between true north and the viewing direction in degrees for each image. Each light grey dot represents one image with a certain average performance level, and the curved black lines indicate the best possible cosine fit of the data. The highest peak of the fitting curve indicates the angle that led to the overall best result.

## Discussion

In the present study, we investigated how spatial information about orientation and location of houses and streets is coded. We performed three experiments requiring different spatial orientation abilities. We investigated the *unitary coding* of objects by testing the knowledge of absolute orientation of single houses and streets towards cardinal directions. Furthermore, we tested *binary coding* by testing relative orientation between pairs of houses and streets and pointing between locations of houses. We distinguished task performance with limited time from performance with infinite time for cognitive reasoning. When spontaneous knowledge retrieval was required, our results show a significantly better performance in assessing the relative orientation of houses to each other (*binary code*, H_1_) than in assessing their individual cardinal orientation (*unitary code*, H_0_). As the *binary code* directly codes object relations, it thus supports spatial behavior and gives direct and fast access to action-related information. This conclusion is supported by our observations from the pointing task, which directly probed action-related knowledge of the direction of an object’s location relative to another object. This task of pointing from one location to another revealed an even better performance. The possibility of using cognitive strategies (infinite response time) enabled participants greatly to improve their judgments of houses’ cardinal orientations, but it influenced the performance of the relative orientation task much less. In contrast to house stimuli, performing the absolute and relative orientation tasks under time pressure with streets as stimuli led to the opposite pattern of results. Streets were significantly better remembered with respect to their cardinal orientation, providing evidence that streets are preferentially coded in a *unitary code* (H_2_). In summary, when spontaneous retrieval is needed, houses are preferentially coded in a *binary code*, which relates two houses to each other. For the *unitary coding* of a single house towards true north, time for
cognitive reasoning is required. By contrast, when spontaneous retrieval is required, streets are preferentially coded in absolute coordinates (in a *unitary code*) directly providing access to directional information.

As former studies investigating spatial cognition abilities found that males reacted faster and performed better in a variety of spatial tasks (Masters and Sanders 1993; Moffat et al. 1998; Newhouse et al. 2007; Woolley et al. 2010), we investigated the influence of gender in our experiments. In line with these previous results, we found that males outperformed females in all three experiments. However, in none of our task results did we find a significant interaction including the factor of gender. Taken together, while in general males performed better than females, the relative performance between tasks was comparable for both genders, and the conclusions presented in this study hold for both genders.

Investigating the influence of actual viewing direction and the orientation of a photograph, Sholl and colleagues (2006, 2008) showed in an allocentric-heading recall task in which the alignment of the participant’s current physical heading and the perspective of a given photograph improved decision speed. These findings were supported by several researchers (Burte and Hegarty 2012, 2014) and are in line with our results from the absolute orientation task. Here, participants performed best when the correct north arrow was pointing right in front of them and thus aligned with their actual viewing perspective. Performance decreased (circularly) with increased angular distance (Frankenstein et al. 2012). Even though this effect was visible with infinite time for cognitive reasoning as well as with restricted time, the alignment effect was considerably more marked with the restricted response time. Previous research found that an alignment between learned and tested viewpoint also improved scene recognition (Diwadkar and McNamara 1997) and pointing accuracy (Shelton and McNamara 1997) as well as increased performance when the imagined body orientation was aligned with the actual body orientation (Kelly et al. 2007). An increased pointing accuracy was also found when imagined viewpoints were aligned with a learned axis of an object array (Mou and McNamara 2002). Interestingly, we found no influence of the current viewing direction on to performance in the relative orientation task, where judgments of house-to-house or street-to-street relations were required. Therefore, our results suggest that in contrast to the absolute orientation task, where the current egocentric viewing perspective has an influence on the performance, the relative orientation task has no such dependency.

In our study, we have to consider that investigating real-world stimuli has advantages and disadvantages. As an advantage, with our study design we directly gain insight into aspects of real-world navigation. However, prior experience with the set of real-world stimuli tested in our experiments might have differed between participants, leading to a difference in familiarity. To ensure some similarity with respect to preexisting experiences, we only included participants who had lived at least one full year in Osnabrück. To be able to investigate the spatial navigation in an urban environment and thus to investigate our experiments on the basis of executed navigation, a future study using a virtual-reality city should be conducted. In such a setup, experience gathered at different locations could be objectively tracked and evaluated. This would address the problems posed by a natural city environment. With the fast development of technologies, such an approach seems to be within reach.

Spatial memory can be acquired by experience, which leads to the formation of local reference frames, and also from maps, which lead to global reference frames (Frankenstein et al. 2012; McNamara et al. 2008; Meilinger 2008b; Meilinger et al. 2014, 2015; Richardson et al. 1999; Sun et al. 2004; Wang and Spelke 2002). For example, a pointing task in a virtual-reality replica of Tübingen revealed that participants used a mental map of the city that was oriented toward north (Frankenstein et al. 2012) but used local reference frames that were encoded parallel to the orientation of local streets when estimating the route to a goal (Meilinger et al. 2013). Local and global reference frames may be combined in an environmental space to refer to a required goal (Meilinger et al. 2014). Combining local reference frames with an oriented global reference frame is advantageous because it permits faster spatial orientation. However, learning spatial relations from experience includes inaccuracies. For example, survey tasks using global reference frames derived from experience yield poorer performance with increasing distance between targets; this is known as the distance effect (Loomis et al. 1993; Meilinger et al. 2015). By contrast, deriving a global reference frame from a map should not lead to such distance errors (Frankenstein et al. 2012). The distance between houses tested in our experiments had a range of up to 2.5 km. Analyzing the distance between tested houses revealed no significant difference in performance in the considered range in our experiments. Importantly, as our design contained not only direct relations between two houses but also triangulations, it was not suitable to investigate a distance effect for paired house relations. Due to the lack of a distance effect, our experiments give no direct evidence for a lack of distance effect and thus do not allow us to distinguish conclusively between a global reference frame learned from experience or from a map.

The concept of spatial reference frames has been widely studied in recent decades (e.g. Goeke et al., 2013, 2015; Gramann 2013; Klatzky 1998; Riecke et al. 2007). It was shown that egocentric and allocentric reference frames are combined when navigating in a natural environment. While learning object locations in a new environment requires active experience in egocentric terms (McNamara et al. 2003), spatial relations may also be remembered in an allocentric reference frame (Mou and McNamara 2002). With the introduction of the relative orientation task, we tested the recall of spatial relations between pairs of houses and streets, whereas the absolute orientation task tested the remembered orientation of single houses and streets with respect to cardinal directions. Our results showed that under time pressure, relative house-to-house orientation and location yielded better performance than individual house orientation to cardinal directions, which supports the hypothesis that spatial knowledge is learned and stored in an action-oriented way. This is in line with previous investigations, which defined an intrinsic reference frame describing object-to-object spatial relations (Mou and McNamara 2002). Remembered allocentric spatial relations are retrieved and combined with present egocentric spatial relations for action in the environment (Kelly et al. 2007; Mou et al. 2004). The assumption that enduring memories of object locations in the environment are represented at least in part in terms of allocentric reference frames is supported by other researchers (Mou et al. 2006; Sargent et al. 2008; Waller and Hodgson 2006). In conclusion, our results suggest that remembered allocentric spatial relations can be coded in an action-oriented way.

Egocentric and allocentric spatial reference frames play an important role in spatial navigation and action selection in general. But the available objects also allow a certain set of actions and, thus, have an influence on the performed action. Forty years back, Gibson (1977) introduced his influential theory on affordances. He put forward that an object has certain action possibilities that an agent can perceive and interact with. Previous research on affordances has mainly dealt with object affordances (Tucker and Ellis 1998). More recently, action possibilities of spatial scenes, especially navigation possibilities, have also been investigated (Bonner and Epstein 2017; Greene and Oliva 2009). Bonner and Epstein (2017) found that action possibilities in an environmental space were automatically extracted, involving the activation of a specific area in the visual center of the human known as the occipital place area. As the two main components of a town, houses and streets, relate to different repertoires of actions, we investigated whether houses and streets would also lead to different results in our experiments. We argue that a single street relates to meaningful actions like locomotion. With houses, the situation appears to be more diverse. Some actions relate to single houses; others, like navigation, relate more to pairs of houses. Our results indicated that participants recalled house orientations better when relative to other houses, whereas street orientations were primarily recalled in relation to cardinal directions. Thus, our results might indicate that different affordances of houses and streets also lead to different types of coding.

Active navigation within the environment is also accompanied by changes in the sensory signals received by an agent. O’Regan and Noë (2001) brought forward the paradigm of sensorimotor contingencies (SMCs) within the framework of Enactivism. SMCs are defined as lawful dependencies between actions and associated perceptions that are learned while interacting with an environment. As the rules that are learned differ between modalities, SMCs are modality specific (O’Regan 2011; O’Regan and Noe 2001). Since it was originally formulated, the SMC paradigm has been broadened. Maye and Engel (2012) suggested long-term regularities of action sequences “like driving to work or baking a cake,” which they called “intention-related” SMCs. They developed a computational model of SMCs with the basic idea of regarding “SMCs as multistep, action-conditional probabilities of future sensory observations” (Maye and Engel 2011). This model was tested on a robot that controlled its own locomotion by developing perceptual capabilities and thus enabling successful navigation within a task environment (Engel et al. 2013). This work shows that spatial navigation is guided by learned sensorimotor interactions with an environment. In line with an SMC framework, we argue on the basis of our results that humans learn action-related affordances of objects in terms of their relative orientation and location while walking through an urban environment. This helps to make correct spatial decisions, even under time pressure, as seen in our relative orientation task. Knowing the exact location or orientation of one object in relation to cardinal directions is less relevant for action. Hence, it takes time and requires cognitive reasoning to calculate the cardinal reference frame that was needed to solve the absolute orientation task. Even when, participants are provided with a sensory augmentation device that gives spatial information of the cardinal north direction spatial relations play an important role in participants spatial perception and action (Kaspar et al. 2014; König et al. 2016). In summary, the concept of sensorimotor contingencies constitutes a promising framework for explaining the results of this study.

## Conclusion

The present study investigates the coding of allocentric information of house and street stimuli relative to a cardinal direction as well as relative to houses or streets, respectively. The results suggest that in case of spontaneous retrieval, the participants primarily relied on house-to-house relations, whereas streets were coded in relation to cardinal directions. With unlimited time for cognitive reasoning, cardinal orientation of single houses improved strongly. This pattern of results supports the view that spatial information about houses’ orientations and locations is primarily learned in an action-oriented way, as proposed by an enactive framework for human cognition. Investigating our experiments on the basis of controlled training using a VR city offers interesting topics for future research.

### Compliance with Ethical Standards

This work was funded by the European Union’s Horizon 2020 program, H2020-FETPROACT-2014, SEP: 210141273, ID: 641321 socSMCs. The authors declare no competing financial interests. All procedures performed in studies involving human participants were in accordance with the ethical standards of the institutional and national research committees. Informed consent was obtained from all individual participants included in the study.

## Acknowledgements

Most of all, we would like to thank all the people who helped with recording and preparing the stimuli. Especially, we thank Antonia Kaiser and Annete Aumeistere, who helped a lot with the recordings.

## Author Contributions

C.G., S.U.K, and T.M. wrote the main manuscript. S.U.K, P.K., C.G., and T.M. proofread and iteratively improved the manuscript. S.U.K. and C.G. recorded the experimental data. C.G. implemented the study and analyzed the data. P.K. suggested the study design and procedures for analysis and supervised the study.

